# Cdc42 is required for lymphatic branching, maturation and valve formation during embryonic development

**DOI:** 10.1101/2020.01.28.923847

**Authors:** Yixin Jin, Jian Wang, Yang Liu, Rui Wang, Hailin Chen, Hongjiang Si, Sathish Srinivasan, Binu Tharakan, Shenyuang L. Zhang, Mariappan Muthuchamy, David C. Zawieja, Xu Peng

**Author notes:** Corresponding author: Xu Peng, M.D., Ph.D.

## Abstract

Cdc42, a Ras-related GTPase that regulates the actin cytoskeleton, was recently shown to play an indispensable role in vasculogenesis and endothelial cell (EC) survival. Here, we determined whether Cdc42 also contributes to lymphatic development by generated two different Cdc42 knockout mice lines by crossing Cdc42/flox mice with either vascular endothelial cadherin-Cre (Cdh5α-Cre) mice or Prox1-CreER^T2^ mice. Our results demonstrated that depleting ECs of Cdc42 expression resulted in embryonic lethality with severe edema. Whole-mount immunofluorescence staining of both knockout embryos showed that the deletion of Cdc42 in ECs impaired lymphatic vessel branching and that the lymphatic lumen size significantly increased. Moreover, we found that the inactivation of Cdc42 compromised mesenteric collecting lymphatic vessel maturation and prevented valve formation. We go on to show that Cdc42 can be activated by vascular endothelial growth factor c (VEGFc) in cultured human dermal lymphatic ECs (HDLECs), and that knocking down Cdc42 expression in these cells using siRNA decreased VEGFc-induced focal adhesion kinase (FAK) phosphorylation. These findings, when taken together with the fact that inactivating FAK plus one allele of Cdc42 in ECs was sufficient to recapitulate the phenotypes of the Cdc42 EC knockout embryos, suggests that Cdc42 and FAK interact genetically during lymphatic development. In addition, Cdc42 and FAK regulated signal transduction is essential for blood and lymphatic vessel separation. Taken together, our data highlights the important role played by Cdc42 in the development of the lymphatic system.

## Introduction

The lymphatic system is a blind-ended network of tubules that plays crucial roles in maintaining tissue fluid hemostasis, orchestrating immune surveillance, and transporting dietary fat from the intestine to the blood (1–3). The lymphatic vascular network is composed of plexuses of lymph capillaries, precollectors, muscularized prenodal collecting lymphatics, and postnodal major lymph conduits such as the thoracic duct (4). To function properly, the lymphatic system must develop its unique architecture in regionally specific and coordinated fashion. At approximately embryonic day 9.5 (E9.5) in the mouse, SRY-related HMG-box (Sox) 18 and COUP transcription factor 2 (COUP-TFII) stimulate a subset of cardinal vein endothelial cells (ECs) to express prospero homeobox protein 1 (Prox1), which specifies the lymphatic EC (LEC) lineage (2, 5, 6). These cells then sprout and migrate out of cardinal veins and intersomitic vessels at multiple sites and form the primary lymph sac and trunk lymphatic vessels. In addition to sprouting from veins, recent studies have reported that LECs can also originate from non-vein sources in the different organs (3) including the heart (7), mesentery (8, 9) and skin (10).

During lymphangiogenesis, LECs must differentiate, migrate, proliferate, and attach to other cells and extracellular matrix in response to various extracellular stimuli (11). Vascular endothelial growth factor c (VEGFc) is one of the most important growth factors that regulates LEC sprouting and migration during lymphatic vessel development (12). Inactivation of VEGFc impairs LEC sprouting from the cardinal vein and disrupts lymphatic sac formation in mice (13). Consistent with these findings, loss of functional VEGFc in patients was shown to cause defective lymphatic function and lymphedema (14, 15).

Transportation of lymph from the periphery organs to the blood relies on functional lymphatic valves and lymphatic muscle cells (4). Lymphatic valves possess two semilunar leaflets that direct lymph movement toward lymph nodes and prevent backflow. In response to extracellular stimulation, the valve-forming cells in the lymphatic vessels adopt an elongated morphology and then condense to form a ring-like structure (16, 17). At the same time, the cells also become clustered and express high levels of Prox1 and forkhead box protein C2 (FoxC2) (17). During the lymphatic valve forming process, planar cell polarity signaling plays an essential role in regulating EC adherent junctions assembly/disassembly (16). However, there is still a good deal to learn regarding the mechanisms involved in lymphatic valve formation.

Cdc42 is a member of the Rho family of small GTPases and functions as a molecular switch that regulates several cellular processes including the formation of filopodia, the establishment of cell polarity, as well as vesicle trafficking (18, 19). Cdc42’s activity is mainly regulated by three groups of proteins: guanine exchange factors (GEF), GTPase activating protein (GAP) and GDP-dissociation inhibitor (GDI) (20). Cdc42 switch II domain contains Tyr64 site and this tyrosine residue can be phosphorylated by Src (21). Cdc42 phosphorylation enhanced the interaction between Cdc42 and RhoGDI and may affect Cdc42 movement to the specific cellular site (21). Importantly, Cdc42 Tyr64Cys mutation was reported in patient with lymphedema (22). Cdc42 has been shown to be required for neuron, pancreas, lung, intestine, craniofacial, and heart development, as well as T cell homeostasis (23–27). We and others have reported that the deletion of Cdc42 in vascular endothelial cells causes embryonic lethality before E10.5 due to vasculogenesis defects (25, 28). However, because our Cdc42 EC-null mice died before the lymphatics system formed, the role of Cdc42 in this process, as well as in later stages of blood vessel development remains unclear. Thus, two new Cdc42 EC knockout mice lines have been generated by crossing Cdc42/flox mice with either vascular endothelial cadherin-Cre (Cdh5α-Cre) (29), or Prox1-CreER^T2^ mice (30), to determine the role of this small GTPase in blood vessel remodeling and lymphatic development. Our results show that Cdc42 is required for lymphatic formation and blood vessel remodeling. Moreover, we also show that Cdc42 and focal adhesion kinase (FAK) work together in regulating lymphatic development during embryogenesis.

## Results

### Inactivation of Cdc42 in vascular endothelial cells caused embryonic lethality with blood vessel remodeling defects

To investigate the role of Cdc42 in blood vessel remodeling and lymphatic development, we generated a Cdc42 EC knockout mouse line (Cdc42^Cdh5KO^) by crossing Cdc42/flox mice (26) with Cdh5α-Cre mice (29), in which Cre recombinase expression occurs two days later than that of Tie2-Cre used in our earlier study (25, 31). To verify Cdc42 deletion in ECs, we harvested E12.5 Cdc42^flox/flox^; Cdh5α-Cre^-^ (designated controls) (Fig. S1 A, B) and Cdc42^flox/flox^; Cdh5α-Cre^+^ (designated Cdc42^Cdh5KO^) embryos (Fig. S1 C, D) and performed immunofluorescence staining by CD31 (Fig. S1 A, C) and Cdc42 antibodies (Fig. S1 B, D). We found CD31-positive blood vessels in the subcutaneous in both control and Cdc42^Cdh5KO^ embryos. Cdc42 was expressed in vascular ECs in the control (Fig. S1B), but its expression was decreased in Cdc42^Cdh5KO^ (Fig. S1D), suggesting that Cdh5α-Cre can delete Cdc42 in vascular ECs.

Screening of 21 litters (105) weaned pups, revealed that 39 (37%) were Cdc42^flox/flox^ controls pups, 38 (36%) were Cdc42^flow/+;^ Cdh5α-Cre^+^ pups, and 28 (27%) were Cdc42^flox/+^; Cdh5α-Cre^-^ pups. The lack of viable Cdc42^Cdh5KO^ mice suggested that the inactivation of Cdc42 in ECs induced embryonic and/or postnatal lethality. The ratio of viable Cdc42^Cdh5KO^ embryos matched Mendel’s law before E14.5. However, the viable Cdc42^Cdh5KO^ embryo ratio decreased after E14.5, and dead and absorbed Cdc42^Cdh5KO^ embryos were detected. Less than 30% Cdc42^Cdh5KO^ embryos survived beyond E16.5.

To further investigate the role of Cdc42 in blood vessel development, control and Cdc42^Cdh5KO^ embryos between embryonic day (E) 12.5 and E14.5 were harvested. E12.5 control and Cdc42^Cdh5KO^ embryos were indistinguishable upon gross examination, and presented the expected blood vessels in the embryo bodies and yolk sac (Fig. S2 A-D). However, by E13.5, around 60% of the Cdc42^Cdh5KO^ embryos started to show blood vessel formation defects in both the yolk sac and embryo bodies, compared to the controls (Fig. S2 E-H). One day later (i.e. E14.5), these defects were even more obvious, and around 50% of the Cdc42^Cdh5KO^ embryos died (Fig. S2 I-L).

To determine the role of Cdc42 in angiogenesis and blood vessel remodeling, we performed whole-mount staining on E11.5 control and Cdc42^Cdh5KO^ embryos using the EC marker CD105. A hierarchical vascular tree was present in the heads of the control embryos (Fig. 1A). In contrast, 50% of the Cdc42^Cdh5KO^ (3 of 6) embryos had blood vessel remodeling defects, including several abruptly disrupted blood vessels in the head (Fig. 1B). Interestingly, we found that Cdc42^Cdh5KO^ embryos exhibited intersomitic blood vessel branching and capillary formation in the trunk region similar to controls (Fig. 1 C and D). These data suggest that Cdc42 is required for blood vessel remodeling and that the removal of Cdc42 in ECs results in embryonic lethality.

**Figure 1.**
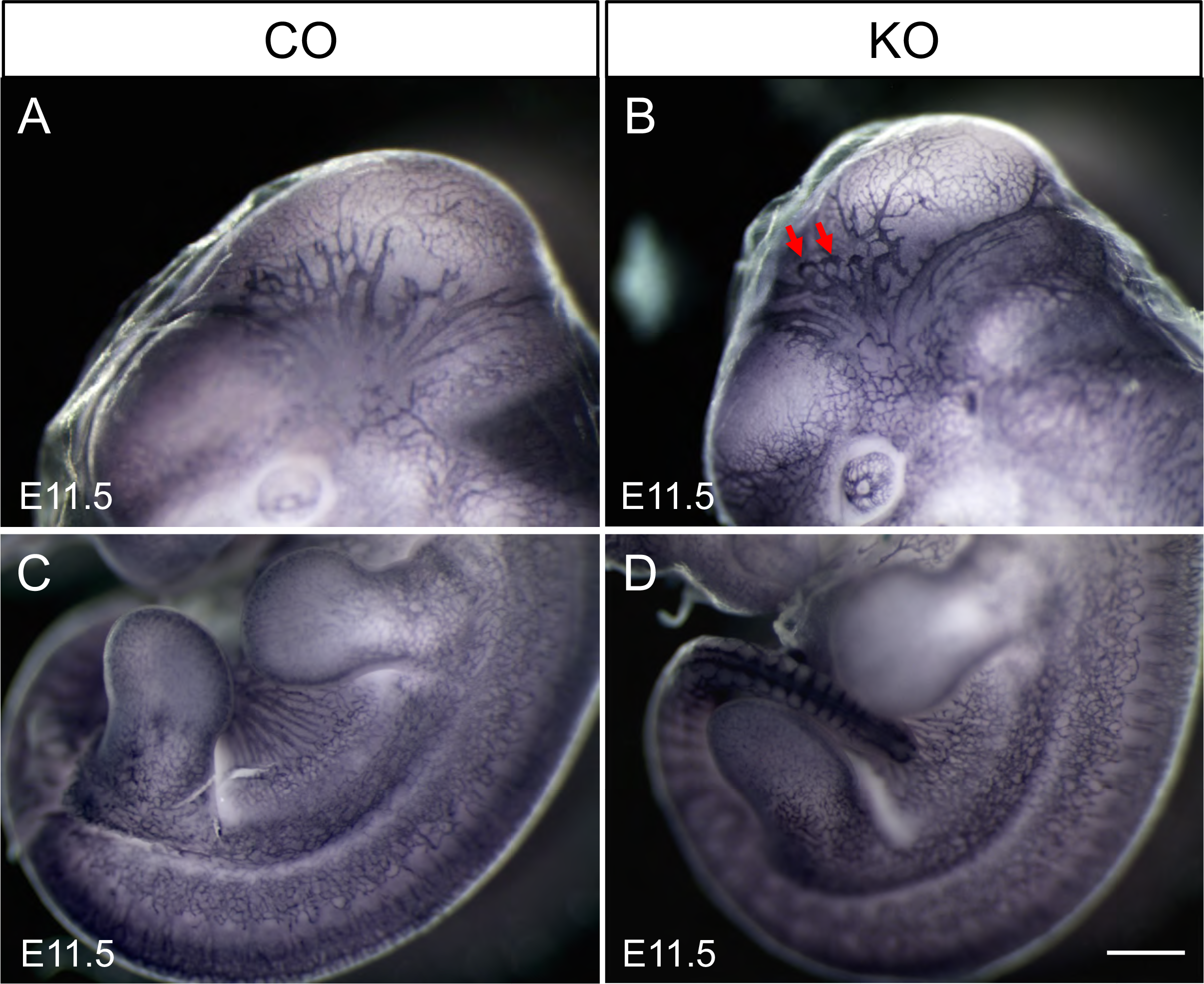
Inactivation of Cdc42 impaired blood vessel remodeling in the brain. Whole-mount staining with CD105 was performed on E11.5 Cdc42^flox/flox^ control (CO) (A, C) and Cdc42^Cdh5KO^ (KO) embryos (B, D). Red arrows indicate abruptly disrupted blood vessels. Bar: 50 μm

### Inactivation of Cdc42 in vascular ECs caused severe edema and lung inflation defects

A small percentage of the Cdc42^Cdh5KO^ embryos survived to birth. However, these pups showed severe edema with purple skin and perished within a couple of hours (Fig. 2A, 2B). The distance between the front lower lip and upper ventral neck of the Cdc42^Cdh5KO^ pups were significantly shorter than that of control animals (Fig. 2A, 2B yellow arrows), indicating severe subcutaneous edema. Functional lymphatic vessels in the lungs is required for lung inflation in new born pups (32). Since the Cdc42^Cdh5KO^ pups that survived to birth appeared purple in color, we next examined whether the deletion of Cdc42 in ECs affected lung inflation. Hematoxylin and Eosin (H & E) staining was performed on P1 control and knockout (Fig. S3 A-D). The alveolar interstitial thickness of the Cdc42^Cdh5KO^ pups (Fig. S3 B, D) was significantly increased compared to control (Fig. S3 A, C), indicating that the deletion of Cdc42 in ECs may interfere with lung inflation due to edema. We also examined viable E18.5 Cdc42^Cdh5KO^ embryos and found profound edema (Fig. 2D), but we did not detect any obvious edema in its littermate controls (Fig. 2C). H&E staining showed the distinct extension in the dorsal subcutaneous tissues in these knockout embryos (Fig. 2E, 2F).

**Figure 2.**
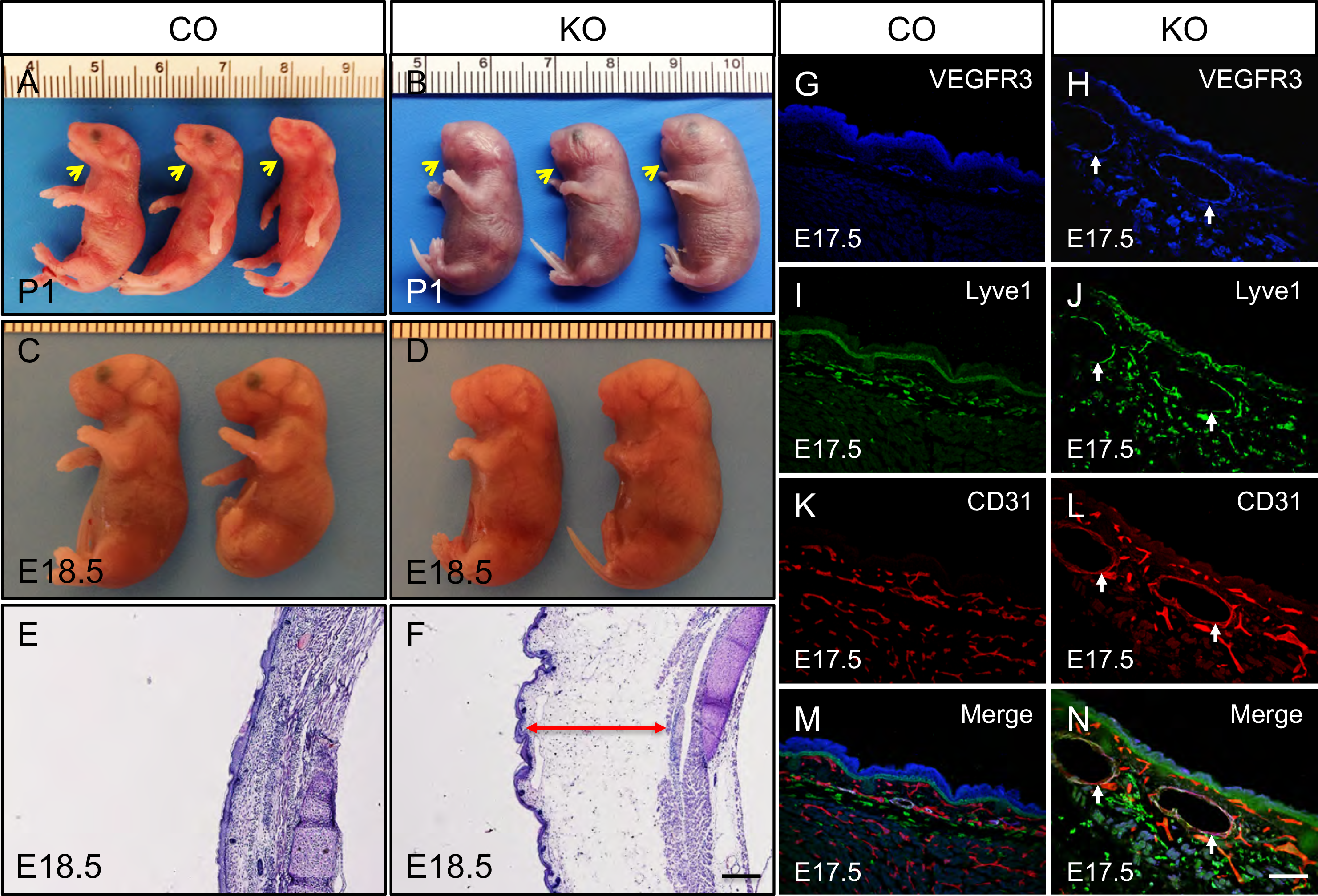
Cdc42 is required for lymphatic development during embryogenesis. Gross examination of P1 pups (A, B) and E18.5 embryos (C, D). P1 Cdc42^Cdh5KO^ (KO) pups showed severe edema with purple colored skin (B), but P1 Cdc42^flox/flox^ control (CO) pups presented no signs of edema with pink colored skin (A). Side view of E18.5 KO embryos showed edema, but no edema displayed in CO embryos (C). Histological analysis (H&E) on the dorsal skin of E18.5 CO (E) and KO embryos (F). The subcutaneous tissue of E18.5 KO embryo exhibited distinct expansion (red line) (F). Immunofluoesence staining with VEGFR3 (G, H), Lyve1 (I, J) and CD31 (K, L) on CO (G, I, K, M) and KO embryons (H, J, L, N). The dilated lymphatic vessels are indicated by white arrows. M and N are merged images. Bars: 125 μm (F), 100 μm (N)

The severe edema observed in the Cdc42^Cdh5KO^ embryos prompted us to examine the effects of deleting Cdc42 in ECs on lymphatic development in the skin. We performed immunofluorescence staining using lymphatic vessel endothelial hyaluronan receptor 1 (Lyve-1), a marker of LECs, on dorsal skin sections taken from E17.5 embryos. The resulting images showed dilated lymphatic vessels in the skin of the E17.5 Cdc42^Cdh5KO^ embryos (Fig. 2H, 2N), but not in the controls (Fig. 2G, 2M). Staining for VEGFR3, which is highly expressed in LECs, showed similar results as the Lyve1 staining (Fig. 2I, J). However, CD31-positive blood vessels did not show any signs of dilation in the Cdc42^Cdh5KO^ embryos, compared to controls (Fig. 2K, 2L).

To further examine the effect of Cdc42 inactivation on lymphatic development, we performed whole-mount staining on E17.5 control and Cdc42^Cdh5KO^ embryonic skins using CD31 and VEGFR3 antibodies. While the CD31 staining showed well-formed blood vessel networks in both the control (Fig. 3A) and Cdc42^Cdh5KO^ embryos (Fig. 3E), the lymphatic lumen size was significantly enlarged in Cdc42^Cdh5KO^ (Fig. 3F, 3G, and 3I, white arrow), compared to controls (Fig. 3B and 3C). High magnification images taken of the samples showed filopodium-like protrusions in the control LEC (Fig. 3D, yellow arrows), but not in the Cdc42^Cdh5KO^ embryos (Fig. 3H). Similarly, E18.5 dorsal skin from Cdc42^Cdh5KO^ embryos stained for Lyve1 and CD31 had dilated lymphatics (Fig. 3J, 3K, yellow arrows). To determine the relationship between lymphatic dilation and edema, we harvested E14.5 Cdc42^Cdh5KO^ embryos with no obvious signs of edema. Whole mount staining of these sections revealed that the inactivation of Cdc42 resulted in lymphatic dilation in Cdc42^Cdh5KO^ embryos (Fig. S4D, 4E, 4F), compared to controls (Fig. S4A, 4B, 4C). Taken together, these data suggest that interrupting lymphatic formation could be a potential reason for the edema observed in Cdc42^Cdh5KO^ embryos.

**Figure 3.**
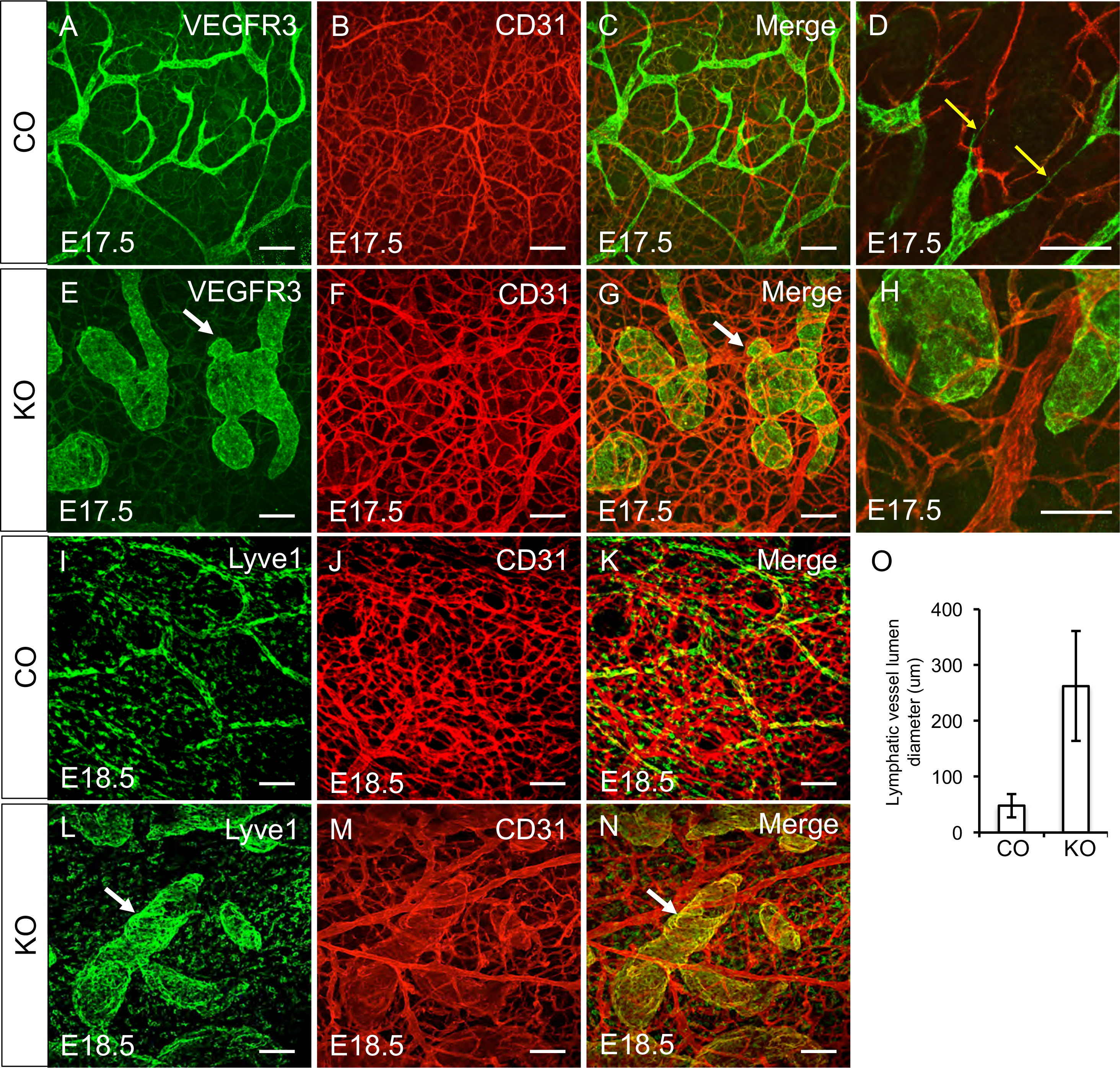
Loss of Cdc42 prevented LEC filopodia formation and increased the lumen size of lymphatic vessels in the skin. Double immunostaining of E17.5 Cdc42^flox/flox^ control (CO) (A-D) and Cdc42^Cdh5KO^ (KO) embryonic skin (E-H) with CD31 (A, D) and VEGFR3 (B, F). C and G are merged images. D and H are magnificent images of the box that show the filopodia (yellow arrows) in the CO (D), but not in the KO (H). (I) Quantification analysis of lymphatic lumen size displays the significant increase in the KO embryos. Double immunostaining of E18.5 CO (I-K) and KO embryonic skin (L-N) with CD31 (I, L) and Lyve1 (J, M). K and N are merged images. The lymphatic lumen size was increased in KO (G, N), compared to the CO (C, K). Bars: 100 μm

### Deletion of Cdc42 in ECs impaired collecting lymphatic maturation in the mesentery

Mesenteric collecting lymphatic is important in transporting absorbed lipids to the blood (4). To determine the role of Cdc42 in this process, we harvested the mesentery from E18.5 (Fig. 4A-F) control and knockout embryos, and then stained them for Lyve1 (Fig. 4A, 4D) and CD31 (Fig. 4B, 4E). The collecting lymphatic vessels of the control mesentery possessed highly branched vessels along the mesenteric blood vessels and displayed relatively even diameters (Fig. 4A). However, Lyve1-positive Cdc42^Cdh5KO^ mesentery exhibited enlarged and irregular collecting lymphatic vessels (Fig. 4D). Unlike control collecting lymphatic vessels, Cdc42^Cdh5KO^ vessels expressed high levels of Lyve1. To determine whether the defects we observed at E18.5 were developmental delays or interferences of lymphatic development, we examined the structure of the collecting lymphatic vessels at P1 (Fig. 4G-L). Strikingly, the morphology of Cdc42^Cdh5KO^ collecting lymphatic vessels showed very broad, flat lymphatic structures with high Lyve1 expression (Fig. 4J) at P1. In contrast, the lymphatic vessels in the control embryos were of similar size and had lower levels of Lyve1 expression (Fig. 4G).

**Figure 4.**
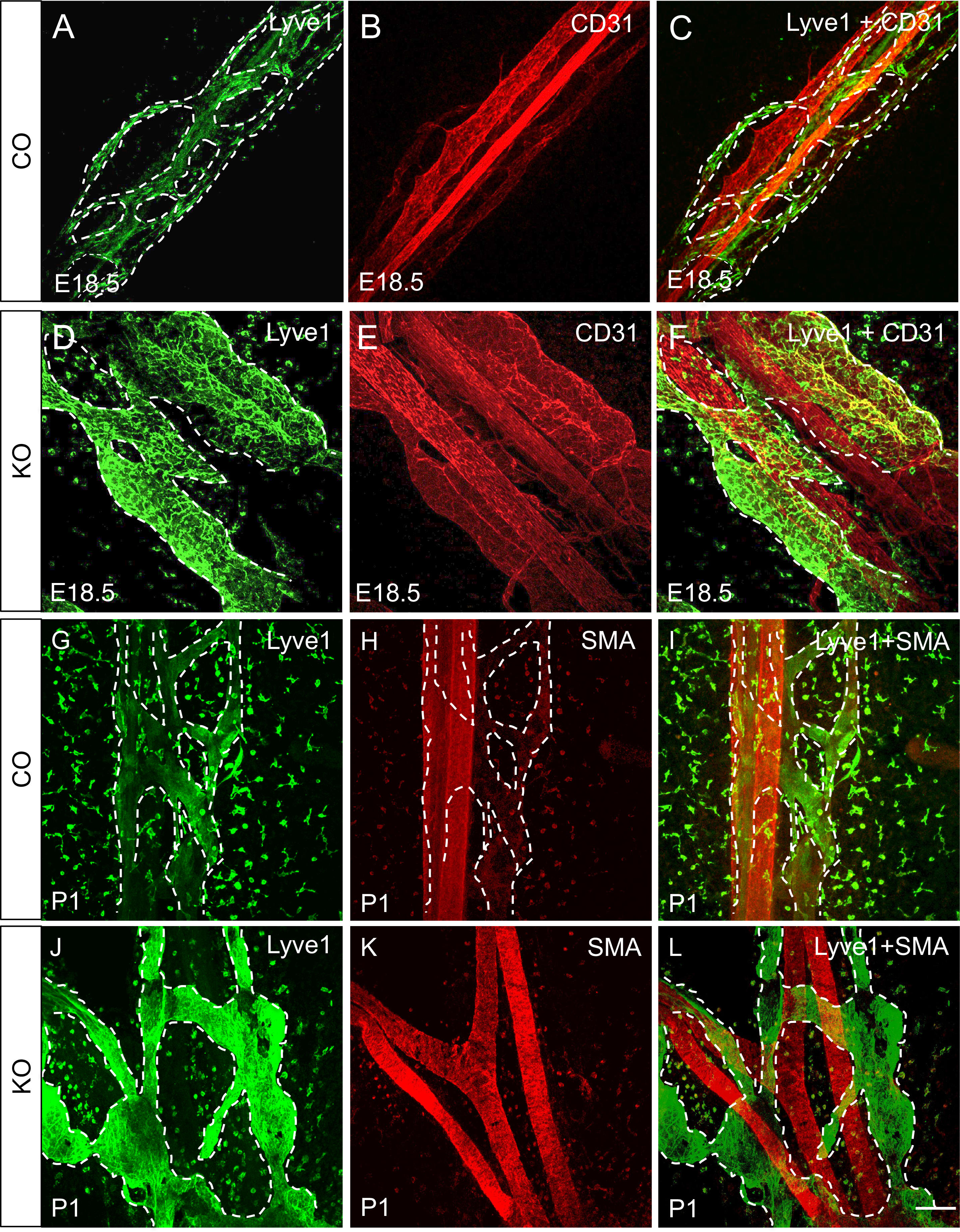
Cdc42 is essential for collecting lymphatic vessel maturation. Immunofluorescence double staining of Lyve1 (A, D) and CD31 (B, E) on mesentery of E18.5 Cdc42^flox/flox^ control (CO) (A, B, C) and Cdc42^Cdh5KO^ (KO) (D, E, F). The lymphatic lumen of KO (F) was larger than that of CO (C). Lyve1 (G, J) and smooth muscle actin (H, K) double staining on P1 CO (G, H, I) and KO mesentery (J, K, L). Smooth muscle actin positive cells were recruited in P1 CO lymphatic (I), but not in KO (L). Bar: 100 μm

Covering the lymphatic collecting vessel with muscle cells is another important step in development. As expected, we found that the collecting lymphatic vessels in P1 control embryos were indeed covered by smooth muscle actin-positive lymphatic muscle cells (Fig. 4H and 4I, white lines and yellow arrows). However, we could not detect any smooth muscle actin-positive lymphatic vessels in the Cdc42^Cdh5KO^ mesentery (Fig. 4K and 4L), indicating that the deletion of Cdc42 impaired lymphatic muscle cell recruitment.

### Cdc42 is required for lymphatic valve formation

Lymphatic valve formation is a sophisticated process that is tightly regulated by the coordinated activation of several different signaling proteins (16, 17, 33). To gain a deeper understanding of how Cdc42 regulates this process, we performed Prox1 and CD31 co-staining on knockout embryos and neonatal pups between E16.5 and P1. At E16.5, the mesenteric lymphatic vessels expressed Prox1 uniformly and the lymphatic valves had not formed yet in either the control (Fig. S5A-S5C) or Cdc42^Cdh5KO^ (Fig. S5D-S5F) embryos. By E17.5, a group of LECs which expressed high levels of Prox1 was observed along the longitudinal axis of lymphatic vessels in both control (Fig. S5G-S5I) and Cdc42^Cdh5KO^ (Fig. S5J-S5L). However, a small cluster of lymphatic valve-forming cells (expressing high levels of Prox1) reoriented 90 degrees and migrated toward the lumen in the control (Fig S5G), but failed to do so in the Cdc42^Cdh5KO^ embryos (Fig. S5J). Moreover, Prox1 expression in the control embryos (Fig. 5A) was decreased in the lymphangion region but maintained at a high level in the lymphatic valve formation zone. Strikingly, no lymphatic valve-forming zone was detected in the Cdc42^Cdh5KO^ embryos (Fig.5D). At E18.5, lymphatic valve-forming cells, which express high levels of Prox1, condensed and formed a ring-like structure in the lymphatic vessels of the control embryos (Fig. 5A). However, the expression level of Prox1 in Cdc42^Cdh5KO^ LECs was even and lacked an obvious ring-like structure (Fig. 5D). In P1 neonatal pups, the lymphatic valve of controls was mature and formed leaflets (Fig. 5G, 5H, and 5I). However, the expression level of Prox1 was evenly high in lymphatics of Cdc42^Cdh5KO^ (Fig. 5J, 5K, and 5L), suggesting that Cdc42 is required for valve formation.

**Figure 5.**
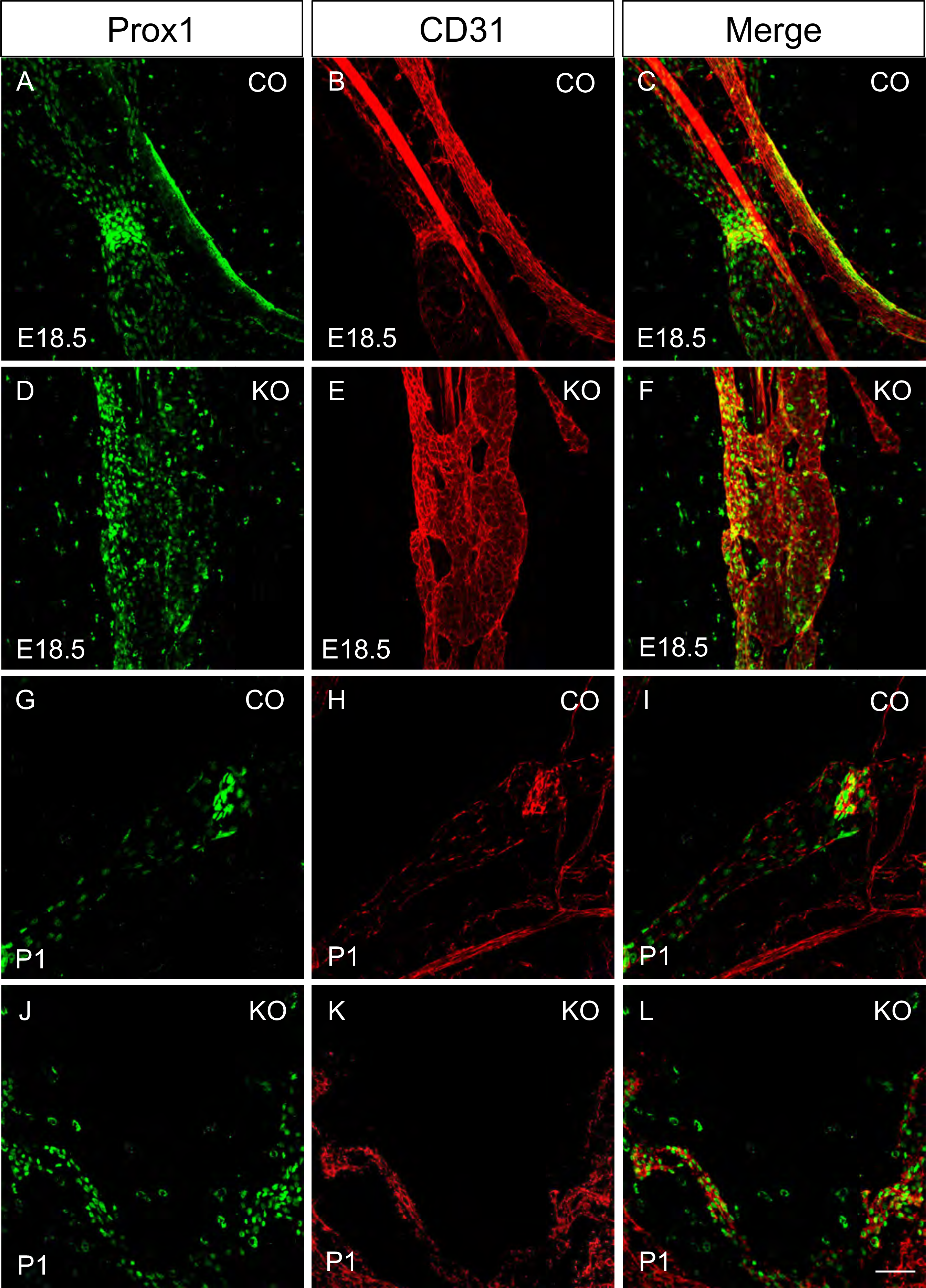
Cdc42 is required for lymphatic valve formation. Immunofluorescence double staining of CD31 (A, D) and Prox1 (B, E) on E18.5 Cdc42^flox/flox^ control (CO) (A, B, C) and Cdc42^Cdh5KO^ (KO) mesentery (D, E, F). Prox1 expression was increased in the lymphatic valve formation region, but expressed in comparable level in the whole lymphatic. P1 mesenteric vessels were stained with CD31 (G, J) and Prox1 (H, K). I and L are merged images. P1 lymphatic valves formed mature leaflets with high Prox1 expression in the CO (I), but no lymphatic valves were evident in the KO (L). Bar: 100 μm

To further confirm the importance of Cdc42 in lymphatic valve formation, we also stained the mesentery collected from P1 knockout embryos with another lymphatic valve marker, integrin α9 (Fig. S6B, S6E), and CD31 (Fig. S6A, S6C). Similar to the Prox1 staining results, V-shaped lymphatic valves were detected in the control embryos (Fig.S6A-S6C), but no lymphatic valves formed in the Cdc42^Cdh5KO^ embryos (Fig. S6D-S6F). Taken together, Cdc42 is essential for lymphatic valve formation.

### The deletion of Cdc42 upregulated VEGFR3 levels in the mesentery collecting lymphatic vessels

The next question that we wanted to address is how deleting Cdc42 from LECs affected the formation of the collecting lymphatic vessel. Previous studies have shown that Cdc42 regulates plasma membrane receptors endocytosis, and since VEGFR3-mediated signal events play an essential role in lymphatic development (12, 34), we investigated whether inactivating Cdc42 influenced VEGFR3 protein levels. Staining E17.5 control (Fig. 6A-6C) and knockout embryos (Fig. 6D-6F) for VEGFR3 expression revealed that VEGFR3 was diffusely expressed in the collecting lymphatic vessels of control embryos (Fig 6A), but was detected as puncta in Cdc42^Cdh5KO^ embryos (Fig. 6D). Consistent with the results from Lyve1 staining, we observed dysmorpholgical collecting lymphatic vessels in P1 Cdc42^Cdh5KO^ (Fig. 6J-6L), compared to the control (Fig. 6G-6I). At P1, we could observe a relative high expression of VEGFR3 in the lymphatic valve of the control (Fig. 6G). However, the VEGFR3 expression level significantly increased in Cdc42^Cdh5KO^, and obvious lymphatic valves could not be detected (Fig. 6J). These data suggest that depleting LECs of Cdc42 causes the deregulation of VEGFR3 homeostasis and the lymphatic collecting vessel developmental defects.

**Figure 6.**
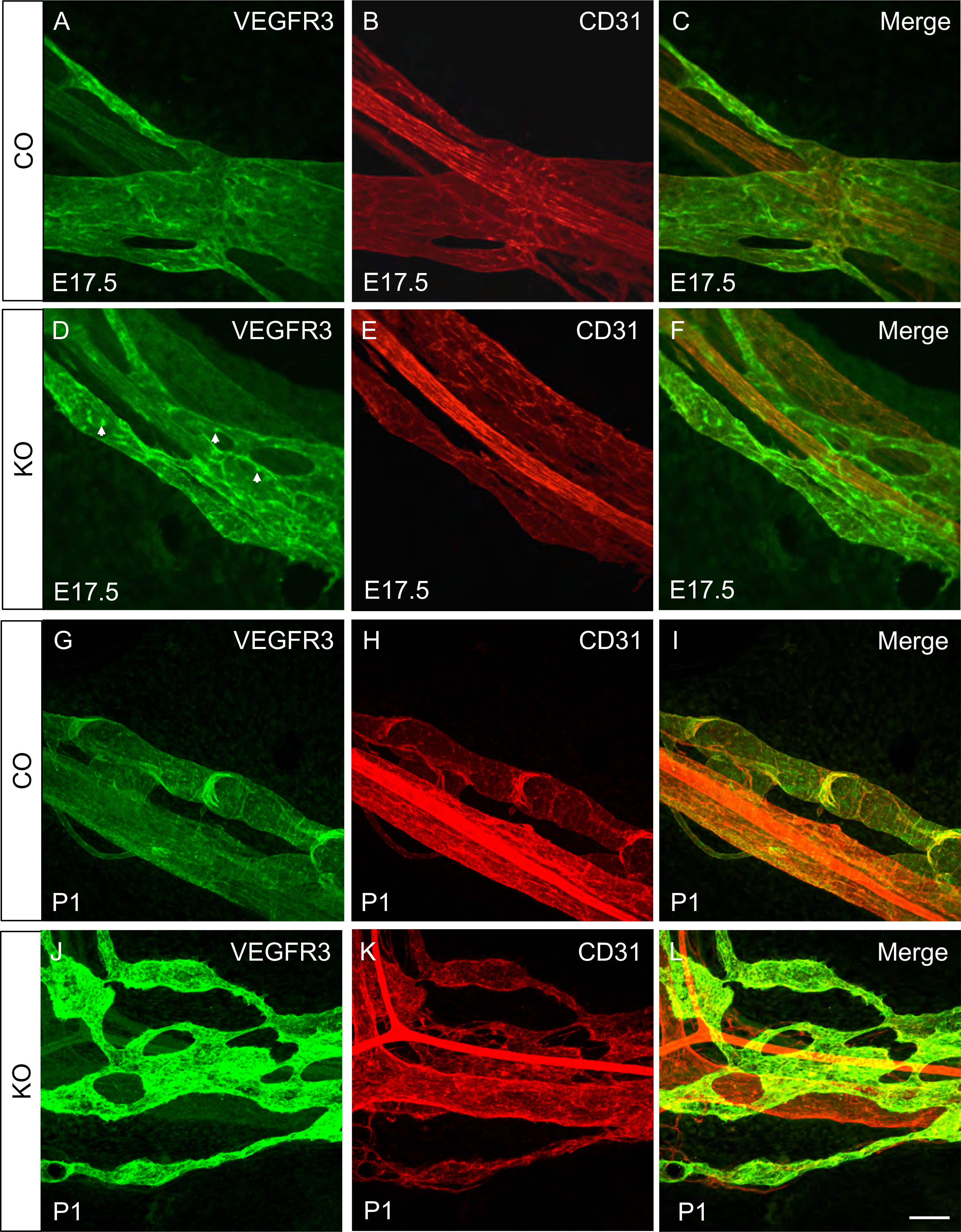
Inactivation of Cdc42 increased VEGFR3 levels in the mesenteric collecting lymphatic vessels. Whole-mount immunofluorescence staining with VEGFR3 (A, D, G, J) and CD31 antibodies (B, E, H, K) on E17.5 (A-F) and E18.5 (G-L) mesentery. VEGFR3 levels slightly increased at E17.5 (D) and dramatically increased at E18.5 (J) in Cdc42^Cdh5KO^ (KO), compared to Cdc42^flox/flox^ control (CO) (A, G). Bar: 100 μm

### The deletion of Cdc42 in LECs resulted in embryonic edema with lymphatic dilation

To further confirm the importance of Cdc42 in lymphatic development, we created a Cdc42 LEC-specific knockout mouse line by crossing Cdc42/flox mice (26) with Prox1-CreER^T2^ mice (30). Our results showed that the inactivation of Cdc42 in LECs resulted in severe edema in E15.5 Cdc42^Prox1KO^ (Cdc42^flox/flox^: Prox1-CreER^T2+^) embryos (Fig. 7A, 7E), and whole-mount dorsal skin staining of these knockout embryos for VEGFR3 (Fig. 8B, 8F) and CD31 (Fig. 7C, 7G) showed dilated lymphatic vessels (Fig. 7F, 7H). In contrast, these characteristics were not observed in the control embryos (Fig. 7B, 7D), suggesting that Cdc42 in LECs is required for lymphatic development during embryonic development.

**Figure 7.**
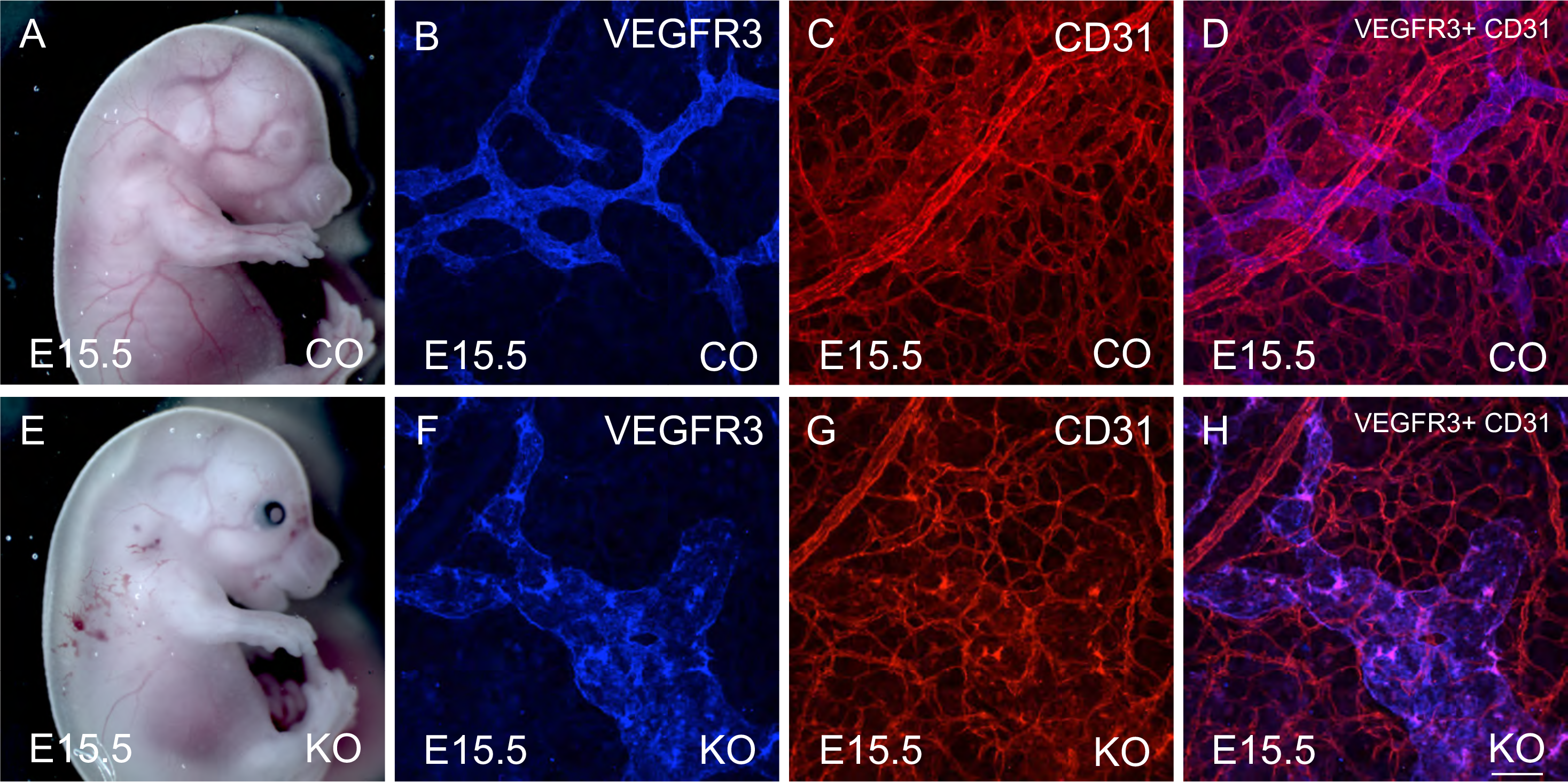
Deletion of Cdc42 in lymphatic endothelial cells resulted in embryonic edema with dilated lymphatic vessels in the skin. Gross examination of E15.5 Cdc42^flox/flox^ control (CO) (A) and Cdc42^Prox1KO^ (KO) (E) embryos. KO embryos displayed edema in the dorsal skin. Double immunostaining of E15.5 CO (B-D) and KO (F-H) embryonic skin with VEGFR3 (B, F) and CD31 (C, G). D and H are merged images. Bar: 100 μm

**Figure 8.**
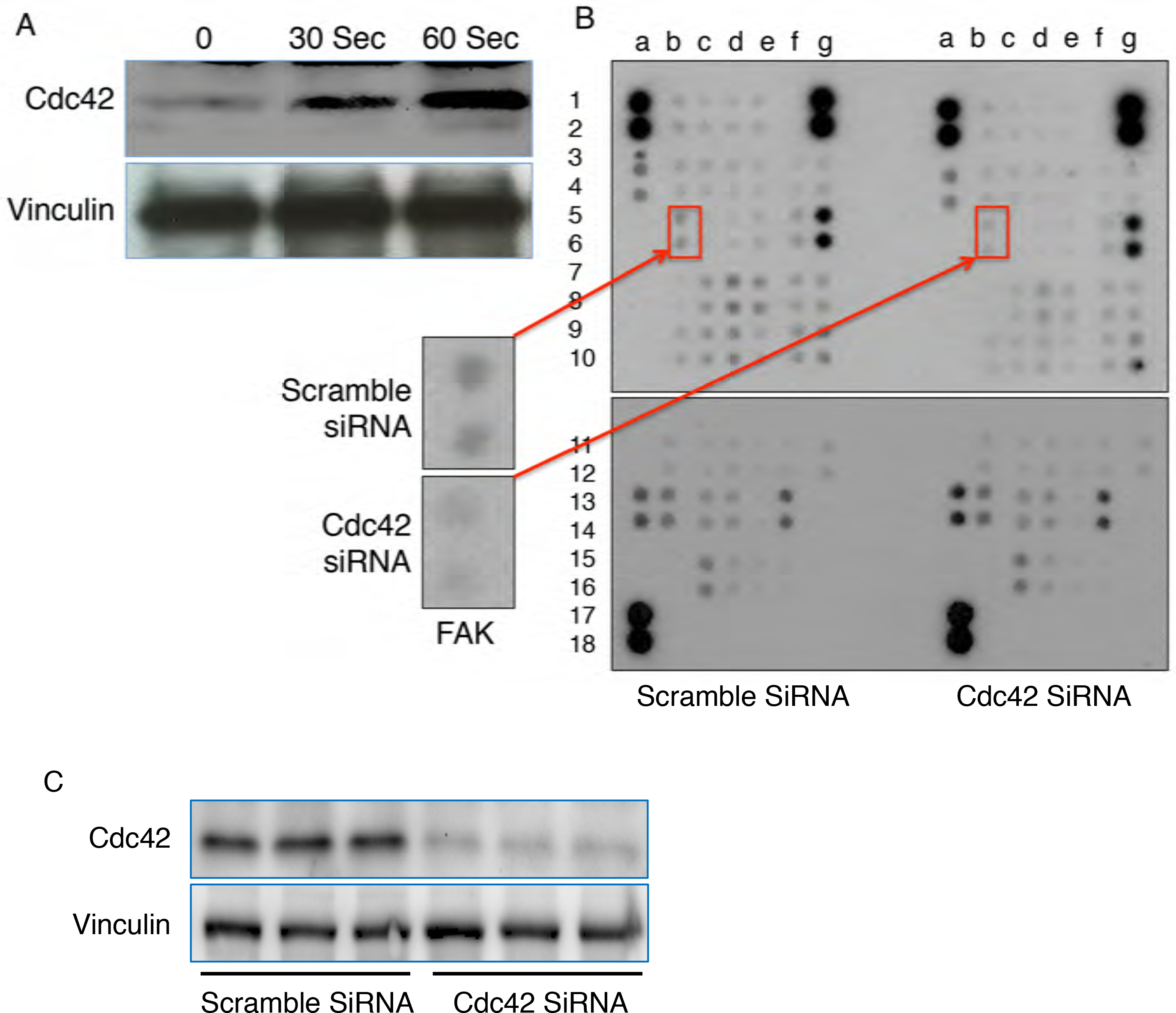
Knocking down Cdc42 in HDLECs impacts VEGFc-induced FAK phosphorylation. (A) HDLECs cell lysates were harvested after VEGFc treatment. PBD pull down assay and Western blots were then performed. The active Cdc42 was detected by Cdc42 antibodies and Vinculin was blotted as a loading control. (B) HDLECs were transfected with scrambled siRNA or Cdc42 siRNA and stimulated with VEGFc. Cell lysates were incubated with Human Phospho-Kinase Array membranes. a1-2, a17-18, g1-2 reference spots. b5, b6 FAK spots.

### Cdc42 is activated by VEGFc and the inactivation of Cdc42 impaired FAK phosphorylation

Cdc42 can regulate multiple cellular processes due to its ability to interact with a variety of effectors. Thus we set out to identify proteins that mediates the ability of Cdc42 to promote lymphatic development is critical for developing novel therapeutic strategies for lymphatic diseases. It is well-known that Cdc42 is activated by the growth factor VEGFa (35–37). Because of the similarities between VEGFa and VEGFc, we investigated whether Cdc42 could also be activated by VEGFc. The human dermal LECs (HDLECs) were treated without or with VEGFc for 30 seconds or 1 minutes. The activated form of Cdc42 in each of the cell extracts was precipitated using the p21 binding domain (PBD) protein, and the resulting complexes were subjected to western blot analysis using Cdc42 antibodies. The resulting blot shows that VEGFc treatment of HDLECs increased Cdc42 activity (Fig. 8A).

We then determined which VEGFc-stimulated signaling events were mediated by Cdc42. For this experiment, cultured HDLECs expressing a scrambled siRNA or an siRNA targeting Cdc42 were treated with VEGFc and lysed. The cell lysates were then incubated with Human Phospho-Kinase Array membranes (Fig. 8B). We found that depleting Cdc42 from these cells did not alter the phosphorylation levels of c-Jun N-terminal kinase (JNK) (a7, a8) or AMP-activated protein kinase (AMPK) (b7, b8). However, FAK phosphorylation (f5, f6) did significantly decreased (Fig. 8B). We and others have previously reported that FAK is important for blood vessel formation (38–41), and since BECs are similar to LECs, it is possible that FAK may mediate the ability of Cdc42 to promote lymphatic development. The aliquot cell lysates were also blotted with Cdc42 and vinculin antibodies to confirm the knockdown efficiency of Cdc42-siRNA (Fig. 8C).

### Cdc42 and FAK coordinated during lymphatic development

To unravel the role of FAK in lymphatic development, we crossed FAK/flox mice with Cdh5α-Cre mice (29) and found that the inactivation of FAK by Cdh5α-Cre did not affect embryonic development (data not shown). However, the inactivation of FAK plus the deletion of one allele of Cdc42 caused severe edema (Fig. 9E), which is similar to the phenotype we observed in Cdc42^Cdh5KO^ pups, but was not observed in the FAK^flox/flox^; Cdc42^flox/+^ control embryos (Fig. 9A).

**Figure 9.**
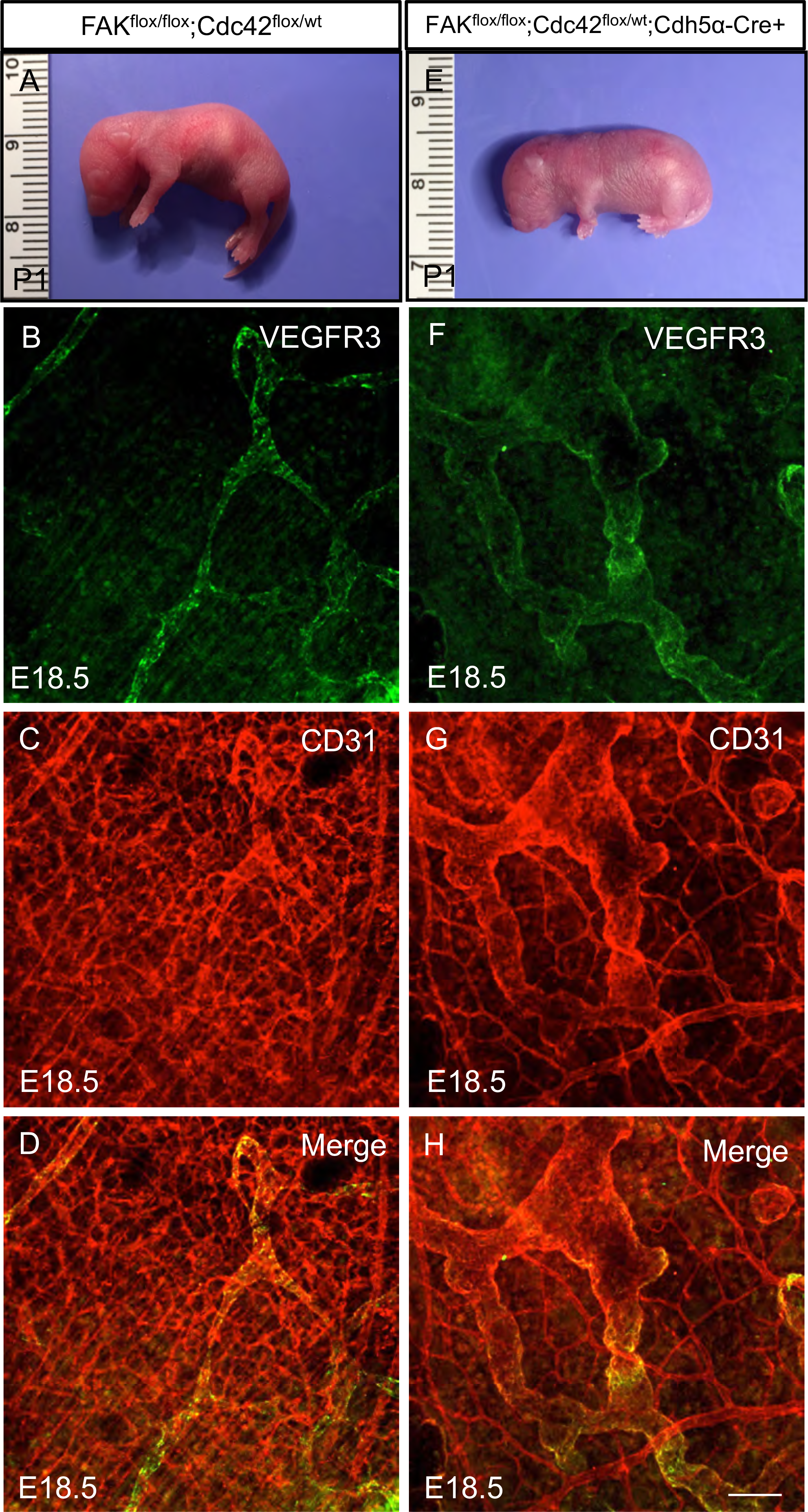
Inactivation of FAK with hypomorphic Cdc42 resulted in profound embryonic edema with dilated lymphatic vessels. Gross examination of E18.5 FAK^flox/flox^; Cdc42^flox/+^ control (A) and FAK knockout plus Cdc42 hypomorphis (FAK^flox/flox^; Cdc42^flox/+;^ Cdh5α-Cre^+^) embryos (B). Whole-mount staining on E18.5 FAK^flox/flox^; Cdc42^flox/+^ control (C, D, E) and FAK knockout plus Cdc42 hypomorphis (F, G, H) embryonic skin with Lyve1 (B, F) and CD31 (C, G) antibodies. Bar: 100 μm

To determine whether there is a genetic interaction between FAK and Cdc42 in lymphatic development, we harvested skin samples from E18.5 control (Fig. 9B-9D) and FAK knockout (with one Cdc42 allele deletion) embryos (Fig. 9F-9H) and stained them with Lyve1 and CD31 antibodies. The images showed that the inactivation of FAK with hypomorphic Cdc42 resulted in subcutaneous lymphatic vessel dilation (Fig. 9F-9H), which mirrored the phenotype observed in the Cdc42^Cdh5KO^ embryos.

The FAK knockout plus Cdc42 hypomorphic embryos also displayed blood-filled lymphatic vessels in the skin (Fig. 10F), but not in the FAK^flox/flox^; Cdc42^flox/+^ controls (Fig.10A). We detected that around 90% of FAK knockout plus Cdc42 hypomorphic embryos showed blood-filled lymphatic vessels. To further examine the blood-filled lymphatic vessels in FAK knockout plus Cdc42 hypomorphic embryos, we peeled the skin off the embryos from FAK knockout plus Cdc42 hypomorphic and FAK^flox/flox^; Cdc42^flox/+^ control embryos. Whole-mount observation showed that there were many blood-filled lymphatic vessels in FAK knockout plus Cdc42 hypomorphic embryos (Fig. 10G). However, we detected very few blood-filled blood vessels in FAK^flox/flox^; Cdc42^flox/+^ control embryonic skin (Fig.10B). Furthermore, we performed whole-mount staining on the skin with CD31 (Fig. 10D, 10I) and Lyve1 (Fig. 10C, 10H) antibodies. We detected spherical lymphatic vessels in FAK deletion plus Cdc42 hypomorphic embryos (Fig. 10H, 10J), but not in the FAK^flox/flox^; Cdc42^flox/+^ control embryos (Fig. 10C, 10E). Interestingly, 3-dimential reconstruction images indicate that the spherical lymphatic vessel is connected to the blood vessel directly (Fig. S7).

**Figure 10.**
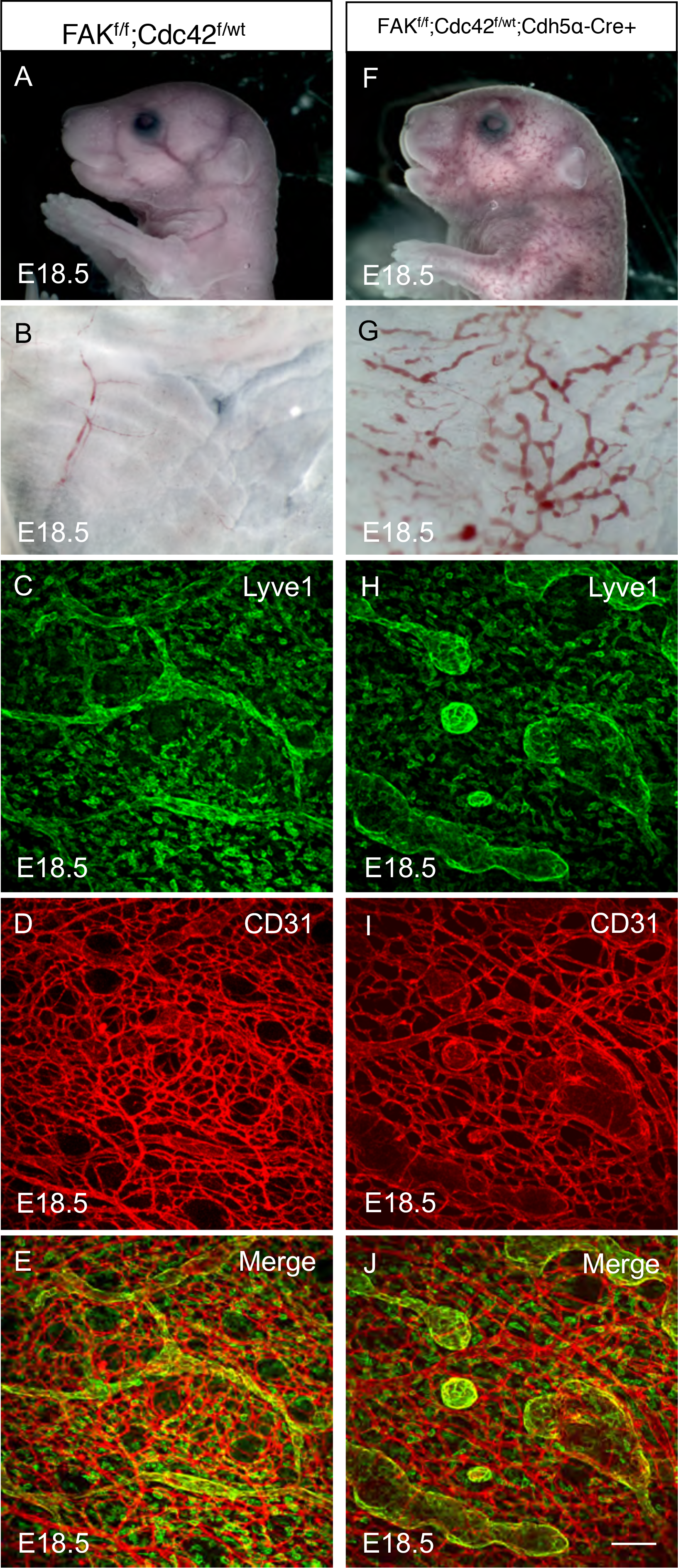
Inactivation of FAK with hypomorphic Cdc42 in ECs results in blood-filled lymphatic vessels. Gross examination of whole pups (A, E) and dorsal skin (B, F) at P1 of control (FAK^flox/flox^; Cdc42^flox/+^) (A, B) and EC FAK knockout with hypomorphic Cdc42 (FAK^flox/flox^; Cdc42^flox/+;^ Cdh5α-Cre^+^) (E, F)). Black arrows indicate blood-filled lymphatic vessels. (C, D, G H). Immunofluorescence double staining of Lyve1 (C, G) and CD31 (D, H) on the skin of E18.5 control (C, D) and FAK knockout with hypomorphic Cdc42 (G, H). White arrows indicate spherical lymphatic vessels that present only in the embryos with blood-filled lymphatic vessels. Bar: 100 μB

## Discussion

During lymphangiogenesis, LECs need to assemble and disassemble cell-cell and cell-matrix adhesions (42). This process is dependent on properly coordinated cytoskeletal remodeling. It is well documented that Cdc42 is an essential regulator of actin polymerization and plays an important role in controlling cell differentiation, migration, proliferation and survival (20). However, the role of Cdc42 in lymphatic vessel formation and maturation remains unknown. In this manuscript, we have demonstrated that the deletion of Cdc42 in ECs leaded to severe edema with dilated lymphatic vessels. Immunofluorescent staining results provided evidence that Cdc42 was required for filopodia formation in the skin. Strikingly, we found that the removal of Cdc42 in ECs prevented lymphatic valve formation, lymphatic muscle cell recruitment, and lymphatic vessel remolding. In addition, we revealed that Cdc42 and FAK were genetically interacted during lymphangiogenesis, and that inactivation FAK plus one allele of Cdc42 caused embryonic edema with dilated lymphatic vessels and defects in blood and lymphatic vessel separation.

As a Rho GTPase family member, Cdc42’s activity is mainly controlled by guanine exchange factors (GEF), GTPase activating proteins (GAP) and GDP-dissociation inhibitors (GDI) (18). Moreover, Cdc42’s functions can also be regulated by tyrosine phosphorylation. Dr. Cerione’s group reported that Tyr64 of Cdc42, which is located in the Switch II domain of Cdc42, can be phosphorylated by Src in response to EGF stimulation (21). Cdc42 Tyr64 phosphorylation enhanced the interaction between Cdc42 and RhoGDI, and Y64A mutation impaired cell transformation (21). Importantly, it was reported that Cdc42 Y64 mutation (Y64C) caused lymph edema in patients, suggesting that defective Cdc42 phosphorylation is important for lymphedema formation (22). Rac1 is another Rho GTPase family member and has an identical amino acid sequence at 61-70 as Cdc42 (43). Interestingly, Rac1 Tyr64 can be phosphorylated by Sr/FAK complex (43). Our data showed that the inactivation of FAK in ECs by Cdh5α-Cre did not result in lymphatic formation defects, but the inactivation of FAK plus one allele of Cdc42 did. This suggests that FAK may work with Src to phosphorylate Cdc42 during lymphatic development. It was well documented that Src/FAK formed a complex to phosphorylate various down steam effectors in different cell types (44). Intriguingly, we also found that the inactivation of Cdc42 impaired VEGFc-induced FAK phosphorylation in cultured HDLECs, but the underlying mechanisms remain elusive. Previously, we reported that the inactivation of Cdc42 downregulated VEGFR2 levels and that VEGFR2 activation can increase FAK phosphorylation (25, 45). Thus, the decreased FAK phosphorylation in Cdc42 knocking-down HDLECs may be due to decreased VEGFR2 levels in LECs.

Filopodia are finger-like membrane protrusions filled with tight F-actin bundles. Filopodia contain growth factor receptors, integrins and Cadherins, etc., and are generally function to environmental probing, cell migration, and cell-cell adhesion formation (46, 47). In response to extracellular stimuli, the active Cdc42 interacts with neuronal Wiskott-Aldrich Syndrome protein (N-WASP)’s GBD/CRIB motif and exposes its C-terminal VSA domain. Subsequently, the active N-WASP induces conformational changes in Actin-Related Proteins (Arp) 2/3, which results in actin polymerization and filopodia formation (48, 49). Alternatively, Cdc42 binds to scaffolding proteins insulin-receptor substrate p53 (IRS53) and interacts with WAVE2 and ENA/VASP to stimulate filopodia formation (50). The first evidence regarding Cdc42 in filopodia formation stemmed from active Cdc42 protein injection-induced filopodia formation in Swiss 3T3 cells (51). Further studies demonstrated that Cdc42 is required in tumor necrosis factor (TNF) α (52), transforming growth factor (TGF) β (53), Neuropilin 1 (Nrp1), and VE-Cadheren-induced filopodia formation (54). However, the deletion of Cdc42 in mouse embryonic fibroblsts did not affect filopodia formation (55), suggesting that the role of Cdc42 in filopodia formation is cell-type dependent. Our data provided solid evidence in favor of Cdc42 being required for LEC filopodia formation. It is well documented that Cdc42 activates atypical protein kinase C (aPKC), and regulates cell polarity establishment and filopodia formation through aPKC-Par6-Par3 complex in various cell types (56). Moreover, aPKC has been demonstrated to play a role in inhibiting stalk EC proliferation during angiogenesis in the retina (57). If lymphatic development shares similar regulatory mechanisms with retina angiogenesis, the deletion of Cdc42 may interfere with aPKC activation and impaired tip lymphatic EC filopodia formation. At the same time, the loss of Cdc42 in the stalk LECs may cause aPKC to lose its inhibitory effect on LEC proliferation, thus causing the dilated lymphatic vessels in the Cdc42 knockout embryos.

The second profound phenotype in our Cdc42 EC knockout mice is the defective collecting lymphatic vessel maturation. We have documented that Cdc42 plays an indispensable role in lymphatic formation and the inactivation of Cdc42 prevented lymphatic valve formation. Interestingly, Cdc42 is not required for the initiation stage of lymphatic valve formation, but is essential for lymphatic valve forming cell condensation to form a ring-like structure. It remains elusive the underlying molecular mechanisms of lymphatic valve forming cell condensation. Differential adhesion hypothesis (DAH) is a hypothesis that explains cell sorting and boundary establishment during embryo development (58). In order to decrease tissue surface tension, the high affinity cells tend to aggregate in the center and the relative lower affinity cells locate in the periphery. It was reported that high prox1 expression LECs tend to aggregate and the expression level of VE-Cadherin in lymphatic valve forming cell is upregulated. In addition, Cdc42 through IQGAP1 enhance cell-cell adhesion. Previous studies have documented that lymphatic valves are formed at the lymphatic vessel connection sites, which face strong turbulence shear stress. Therefore, it is possible that the strong and turbulence shear stress activates Cdc42 and then enhance VE-Cadherin based cell-cell adhesion during lymphatic valve forming cell condensation. When Cdc42 is deleted in LECs, the lymphatic valve forming cells cannot establish strong cell-cell adhesion and impair lymphatic valve forming cell condensation. We also observed that the inactivation of Cdc42 impaired lymphatic muscle cell recruitment. Previous studies have documented that EC and vascular smooth muscles interact through N-cadherin (59, 60), and our studies have demonstrated that Cdc42 is required for N-cadherin-mediated cardiomyocyte adhesions (27). Therefore, it is possible that deleting Cdc42 affects LEC N-cadherin function and results in failed recruitment of lymphatic muscle cells in the mesentery collecting lymphatic vessel.

A recent seminal study showed that VEGFR3 is responsible for the set point response of blood vessels to shear stress (61). We observed that the inactivation of Cdc42 enhanced VEGFR3 expression in the mesenteric collecting lymphatic vessels. It was documented that Cdc42 plays an important role in regulating plasma membrane protein endocytosis (34). Therefore, there is a possibility that the inactivation of Cdc42 impaired VEGFR3 endocytosis and caused VEGFR3 protein levels to be up regulated. Previous studies reported that aPKC-controlled VEGFR3 endocytosis plays a critical role in new blood vessel formation. Our results showed that the inactivation of Cdc42 increased VEGFR3 expression in the collecting lymphatic vessels. It is well documented that Cdc42 can activate aPKC in vitro, so it is possible that the inactivation of Cdc42 decreased aPKC activity and prevented VEGFR3 endocytosis (56, 57). The interfered VEGFR3 endocytosis may cause an increase in VEGFR3-mediated signal transductions, which upregulates Prox1 expression. Another possibility is that the inactivation of Cdc42 disrupted lymphatic valve formation and impaired lymphatic flow. The lack of lymphatic flow may disrupt the down-regulation of Prox1 in the lymphangion. The elevated Prox1 may then enhance VEGFR3 and Lyve1 expression levels. It is also can explain why we only observed VEGFR3 elevation in the mesenteric collecting lymphatic vessels and not in the skin lymphatics.

Deletion of Cdc42 by Cdh5α-Cre caused embryonic lethality after E13.5, which is later than that caused by Cdc42 inactivation by Tie2-Cre (E9.5) (25). Cdh5α-Cre mediated targeted gene deletion occurs as early as E10.5, and the deletion efficiency is around 65% (29). Until E14.5, Cdh5α-Cre deletion efficiency reaches 96%. However, Tie2-Cre mediated gene deletion appears at E8.5 in high efficiency (25, 38). Indeed, this is may be the underlying reason for the difference in embryonic lethality cause between Cdh5α-Cre and Tie2-Cre. Cdh5α-Cre is expressed in all ECs including both LECs and BECs. Therefore, it is possible that both defective lymphatic and blood vessels contribute to the severe edema. We found that the LEC-specific deletion of Cdc42 also resulted in severe edema, suggesting that Cdc42 in LECs plays an indispensable role in edema formation during embryogenesis.

## Acknowledgments

This research was supported by grants from the American Heart Association (19TPA34900011) to X.P.

## Material and Methods

### Generation of Cdc42 EC or LEC specific knockout mice

Cdc42 EC knockout mice were created by crossing Cdc42/flox mice with Cdh5α-Cre mice, and Cdc42 LEC knockout mice were generated by crossing Cdc42/flox mice with Prox1-CreER^T2^ mice. All study protocols were approved by the Institutional Animal Care and Use Committee of Texas A&M Health Science Center.

### Whole-mount staining

The isolated embryos were fixed by the 4% paraformaldehyde (PFA) and dehydrated by methanol. After rehydrated by methanol and PBS, the embryos were incubated with PECAM-1 (1:100) antibodies at 4°C for two overnights followed by HRP-conjugated secondary antibodies. Embryonic skin and mesentery were fixed by PFA for 1 hour at room temperature. The samples were then incubated with different first antibodies, including VEGFR3, LYVE-1, Prox-1, smooth muscle actin, and CD31. After washing 10 mins four times at room temperature, the samples were incubated with the secondary antibodies. The images were photographed using Olympus confocal microscope (Fluoview 300).

### Histological analysis and immunofluorescence staining

The embryos were fixed in 4% PFA for overnight and embedded in Tissue-Tek OCT compound (Electron Microscopy Sciences, PA). After sectioning, Hematoxylin and eosin (H&E) staining were performed to visualize heart and lung morphology. Slides were photographed using an Inverted Olympus microscope (BX51) with a DP71 camera. Immunofluorescence staining analysis was performed as previously descried using different antibodies: VEGFR3, LYVE-1, and CD31.

### siRNA transfection

HDLEC (2 × 10^5^ cells/well) were plated in 6 well plates and incubated with various siRNAs (20 nM) and HiPerFect transfection reagent (Qiagen) for 72 hours, according to the manufacturer’s instructions. Subsequently, HDLECs were used for Western blot.

---

**Figure S1.**
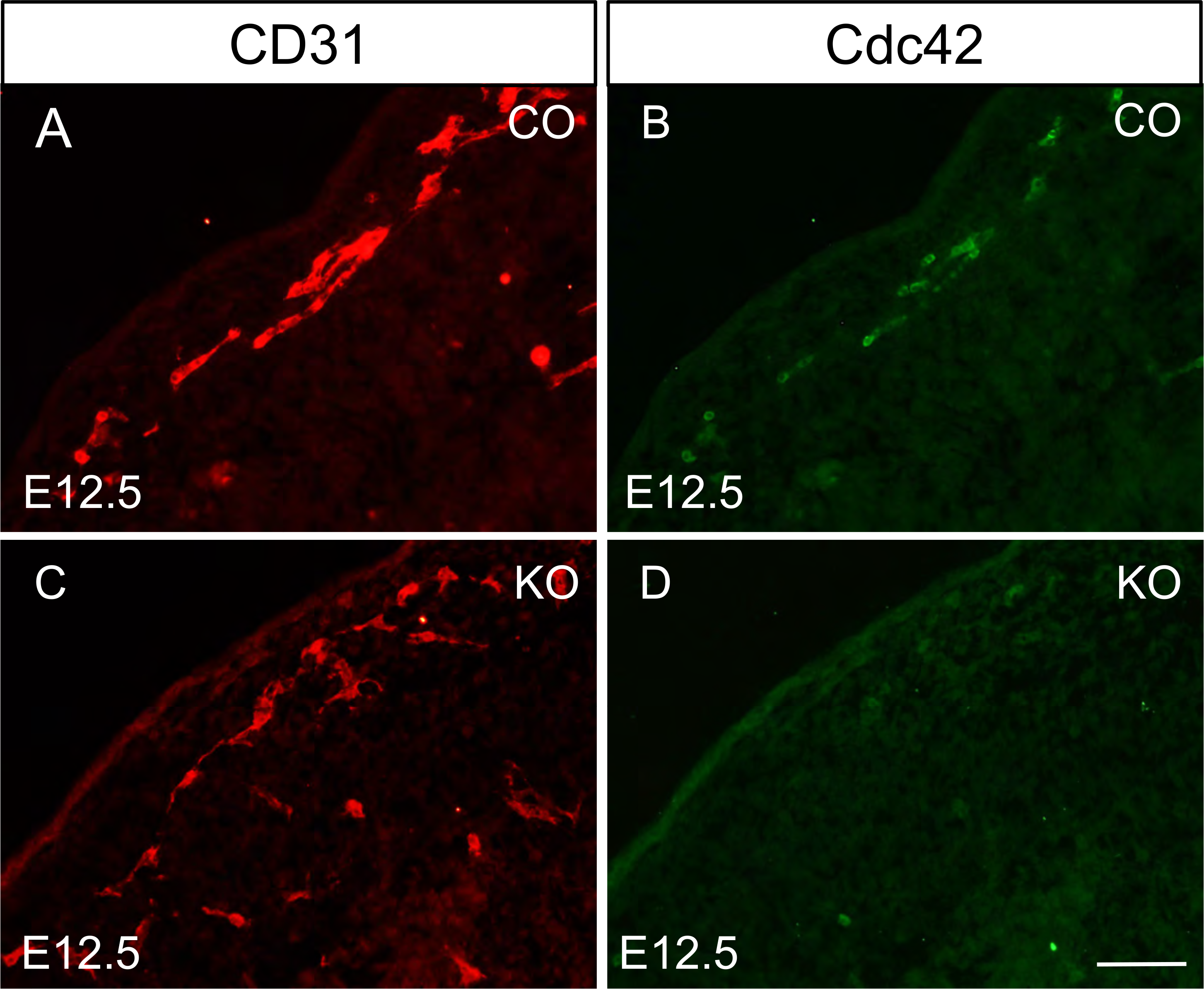
Cdc42 is required for blood vessel formation. Gross examination of Cdc42^flox/flox^ control (CO) (A, B, E, F, I, J) and Cdc42^Cdh5KO^ (KO) (C, D, G, H, K, L) embryos at E12.5 (A-D), E13.5 (E-H) and E14.5 (I-L) with or whitout york sac.

**Figure S2.**
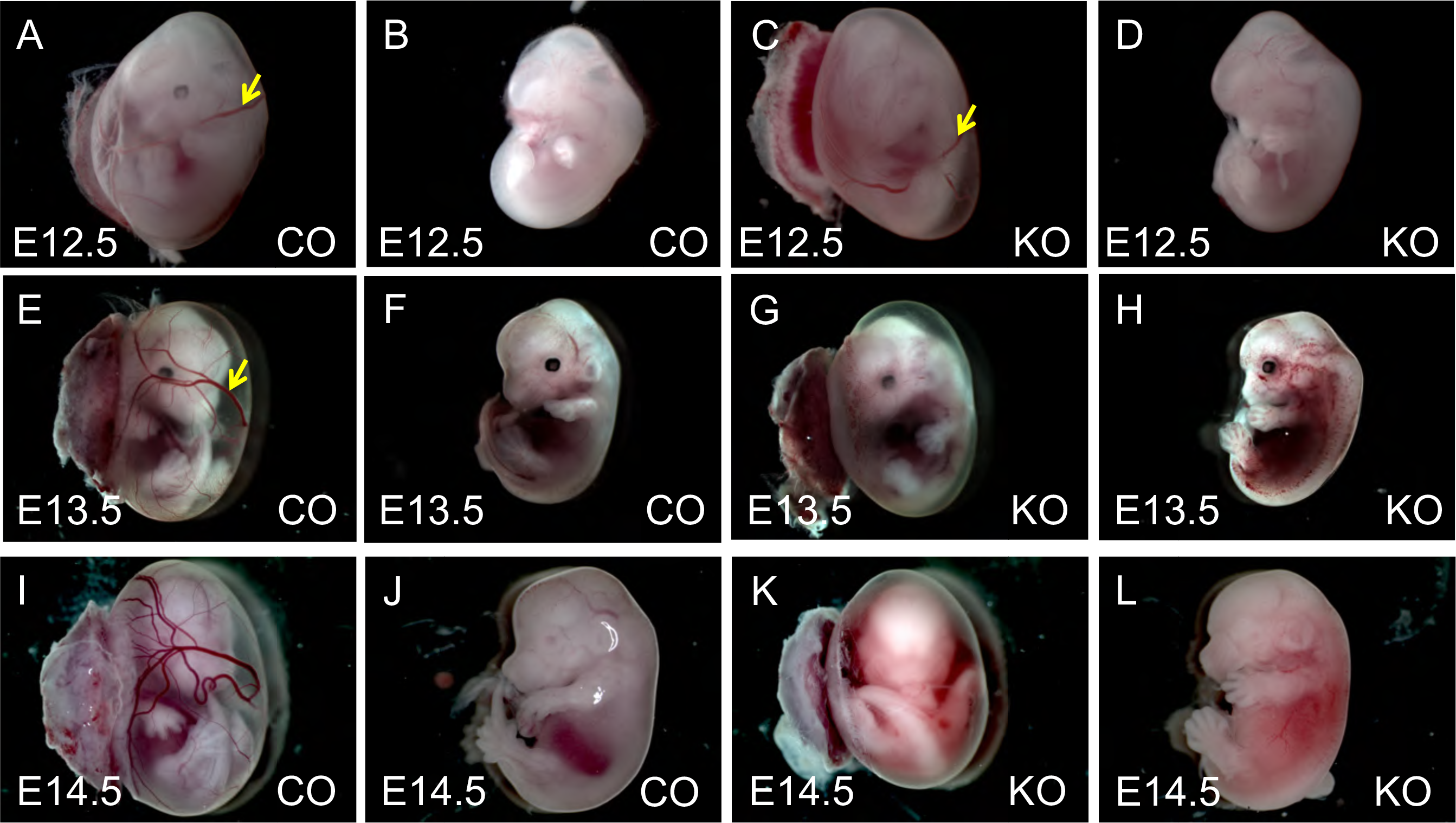
Deletion of Cdc42 in ECs. Sections from E12.5 Cdc42^flox/flox^ control (CO) (A, B) and Cdc42^Cdh5KO^ (KO) embryos (C, D)were stained with CD31 (A, C) and Cdc42 (B, D). Bar: 250 μC

**Figure S3.**
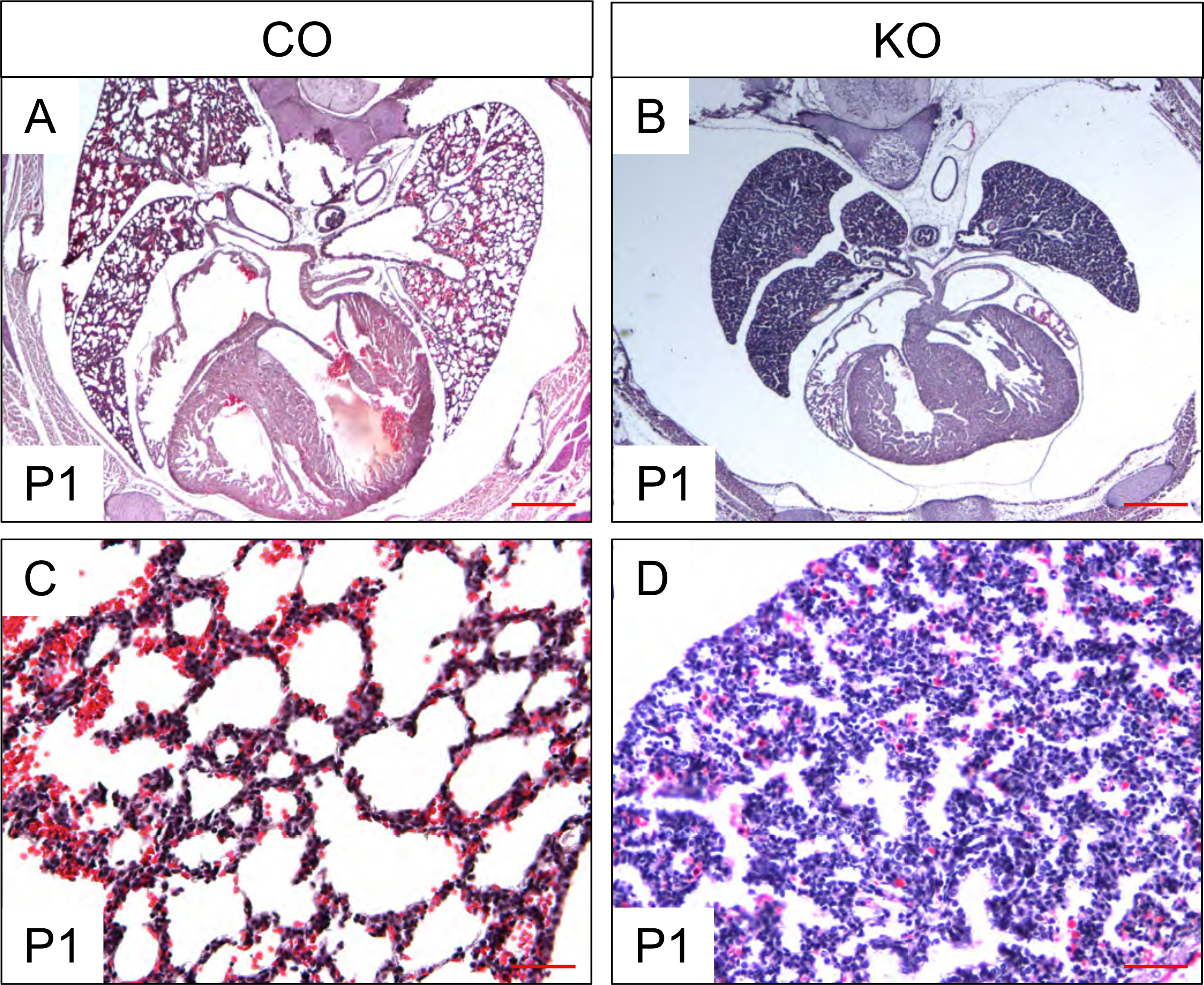
Inactivation of Cdc42 impaired neonatal mice lung inflation. H&E staining anaylsis was performed on P1 Cdc42^flox/flox^ control (CO) (A, C) and Cdc42^Cdh5KO^ (KO) embryos (B, D). (C and D) Higher magnification microphotographs of CO (C) and KO (D). Bars: 500 μK (A, B), 250: 500 CO

**Figure S4.**
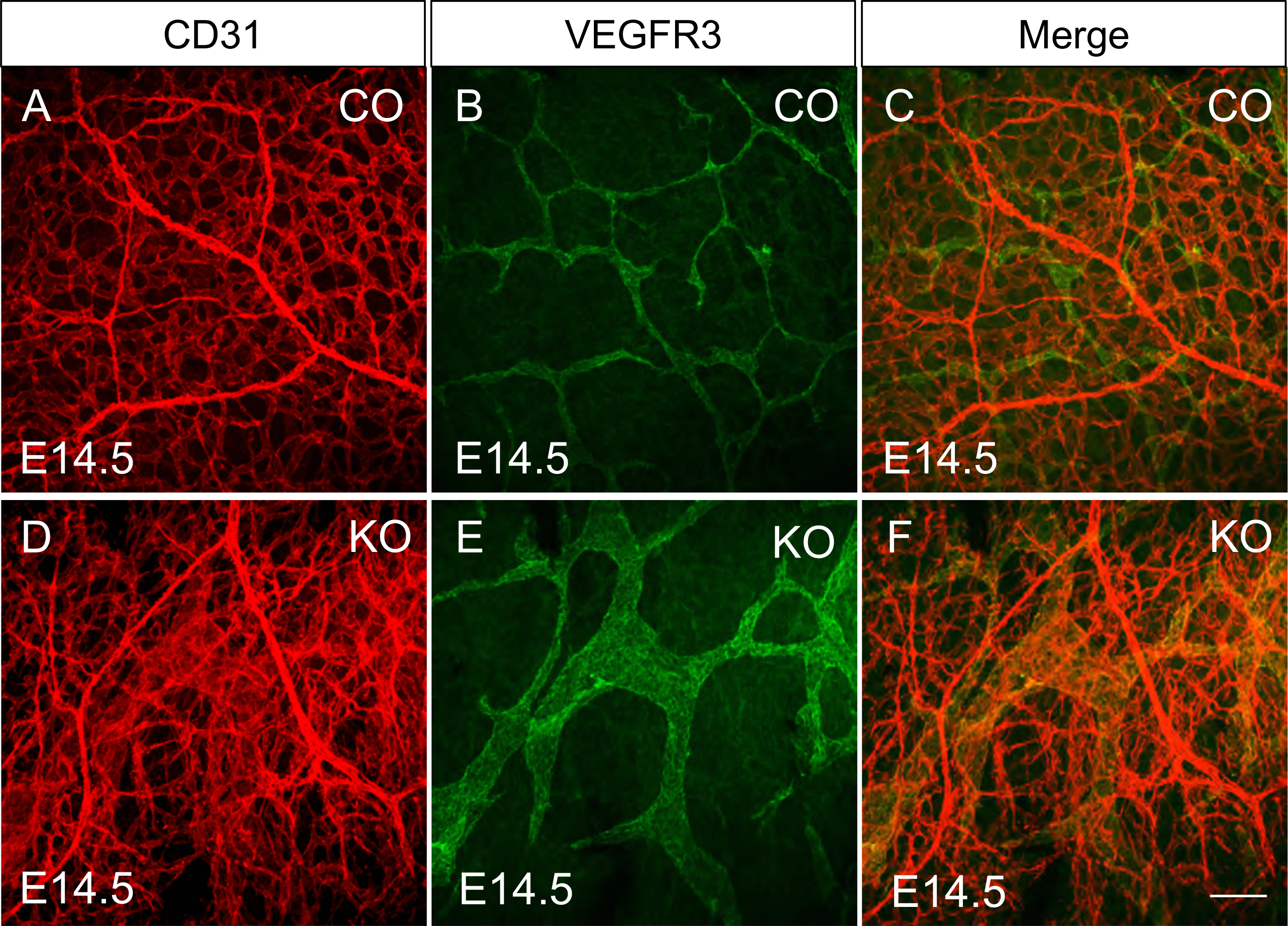
Deletion of Cdc42 resulted in lymphatic vessel dilation. Whole-mount immunofluoresence staining was performed on E14.5 Cdc42^flox/flox^ control (CO) (A, B, C) and Cdc42^Cdh5KO^ (KO) (D, E, F) embryonic skin with CD31 (A, D)and VEGFR3 (B, E). Bar: 100 μV

**Figure S5.**
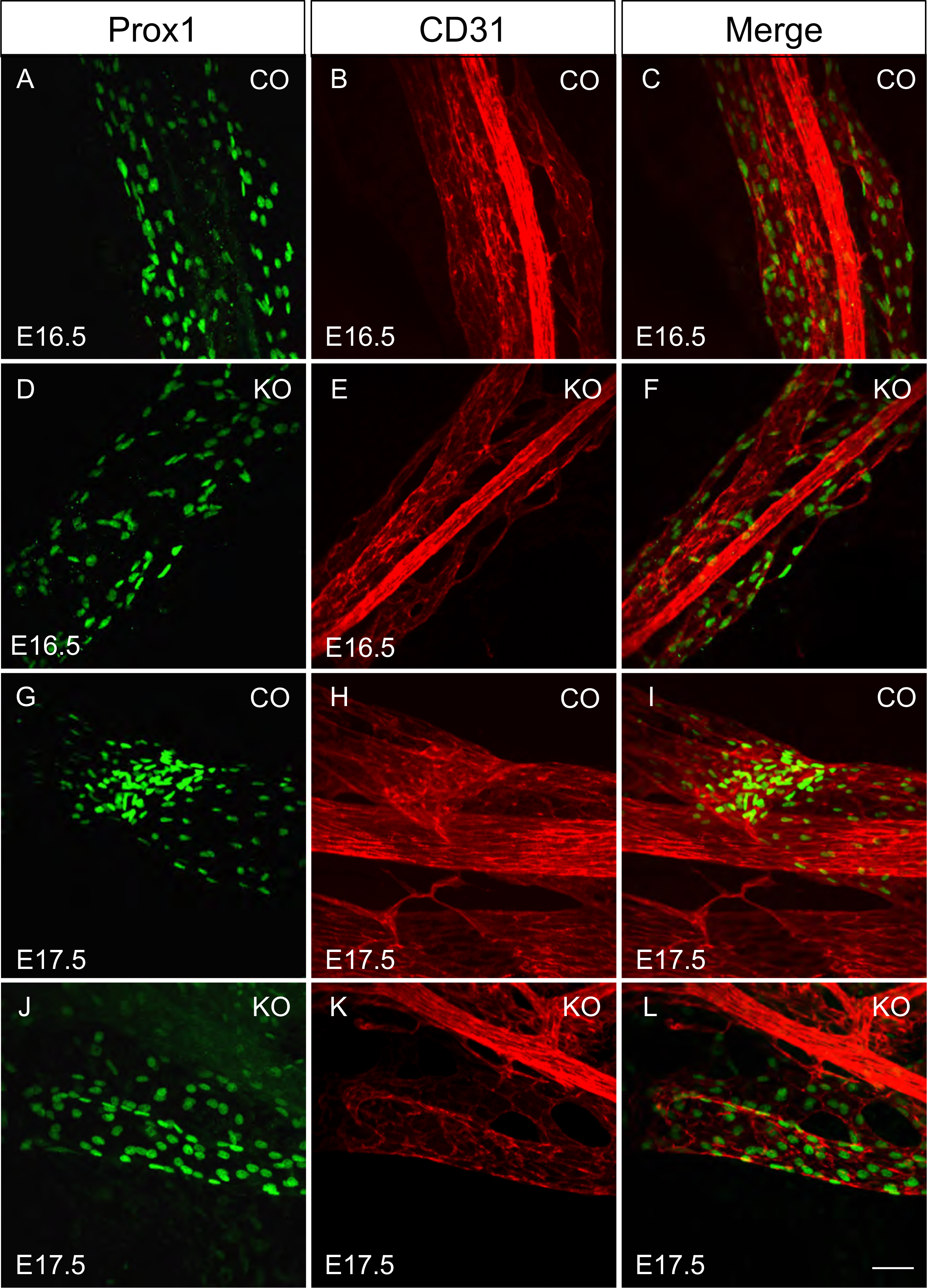
Inactivation of Cdc42 impaired lymphatic valve formaiton. Whole-mount immunofluoresence staining was performed on E16.5 (A-F) and E17.5 (G-L) mesentery with Prox1 (A, D, G, J) and CD31 (B, E, H, K) antibodies. Bar: 100 μa

**Figure S6.**
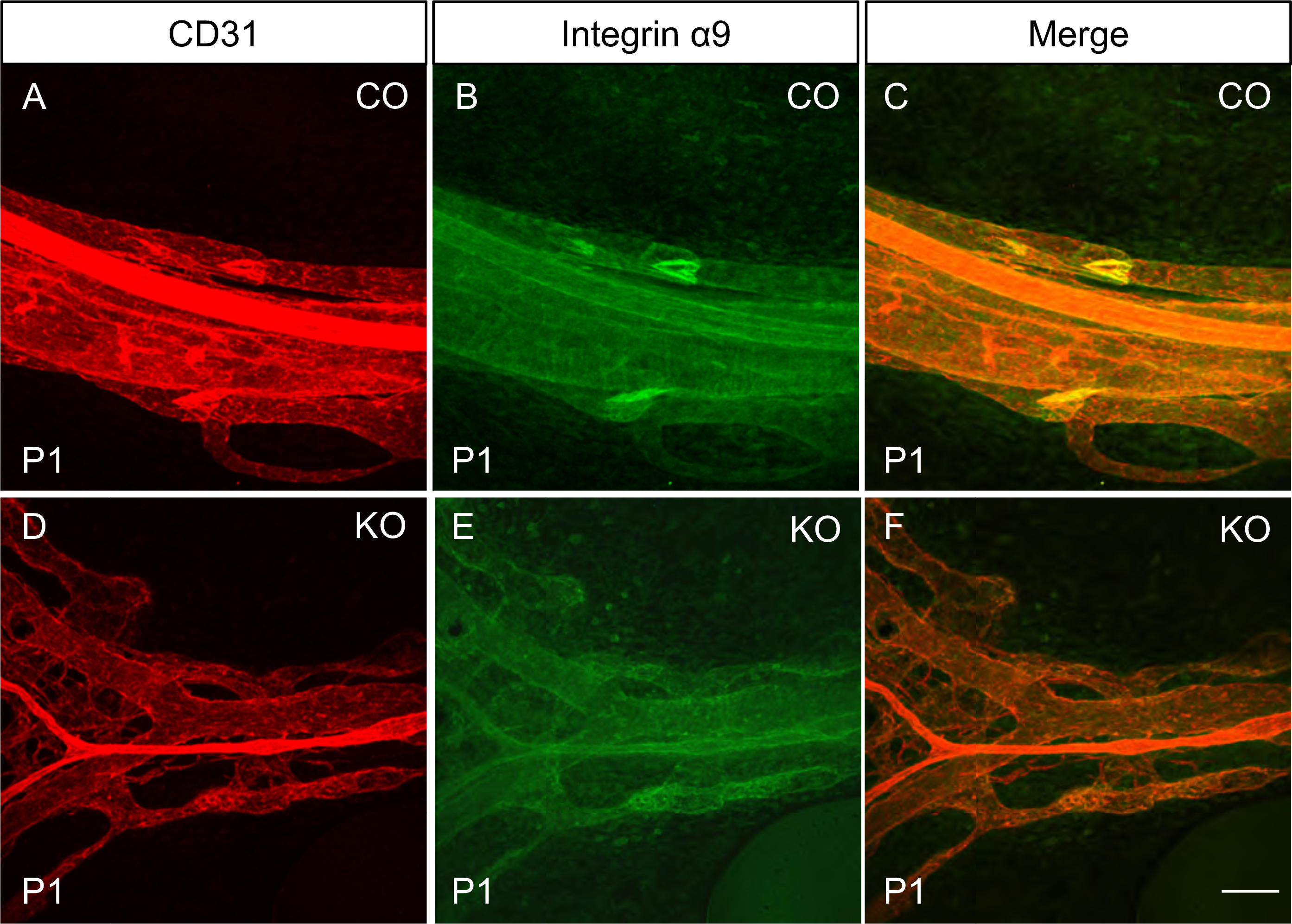
Deletion of Cdc42 prevented lymphatic valve formation. Whole-mount immunofluoresence staining was performed on P1 Cdc42^flox/flox^ control (CO) (A, B, C) and Cdc42^Cdh5KO^ (KO) (D, E, F) mesentery with CD31 (A, D) and integrin α9 (B, E) antibodies. Bar: 100 μB

